# A Methodological Framework to Assess the Accuracy of Virtual Reality Hand-Tracking Systems: A case study with the Oculus Quest 2

**DOI:** 10.1101/2022.02.18.481001

**Authors:** Diar Abdlkarim, Massimiliano Di Luca, Poppy Aves, Sang-Hoon Yeo, R. Chris Miall, Peter Holland, Joseph M. Galea

## Abstract

Optical marker-less hand-tracking systems incorporated into virtual reality (VR) headsets are transforming the ability to assess motor skills, including hand movements, in VR. This promises to have far-reaching implications for the increased applicability of VR across scientific, industrial and clinical settings. However, so far, there is little data regarding the accuracy, delay and overall performance of these types of hand-tracking systems. Here we present a novel methodological framework which can be easily applied to measure these systems’ absolute positional error, temporal delay and finger joint-angle accuracy. We used this framework to evaluate the Meta Quest 2 hand-tracking system. Our results showed an average fingertip positional error of 1.1*cm*, an average finger joint angle error of 9.6^*o*^ and an average temporal delay of 38.0*ms*. Finally, a novel approach was developed to correct for these positional errors based on a lens distortion model. This methodological framework provides a powerful tool to ensure the reliability and validity of data originating from VR-based, marker-less hand-tracking systems.

## 1 Introduction

With recent advances in machine learning, mobile computing and augmented, mixed or virtual reality (commonly known as extended reality or XR), optical marker-less motion-tracking technology has become one of the most cost-effective and easy to implement alternatives for recording hand and finger movements across a wide range of applications, [1–3] This is in contrast to marker-based hand-tracking, here referred to as ground-truth tracking, which requires individual markers to be placed on each tracked finger/hand and a complicated network of specialised and carefully calibrated optical or magnetic sensors, [4–7].

One popular example of this XR marker-less motion-tracking is the Meta Quest 2 (formerly known as Oculus Quest and referred to as Quest, from here on). The Quest hand-tracking system works on the device, without the need for a PC, using a multi-stage process to estimate hand pose and finger angles in real-time, Figure 1, see [4] for more details. The first stage involves correctly identifying the hands from surrounding objects and background (Hand Detection). Next, several key points are identified and labelled on the hand and fingers (Hand Keypoints), which serve as input to an inverse kinematics model of the hand in the next stage (Model-Based Tracking). This uses the key points to estimate global hand pose and finger joint angles. In the final stage, the output of this model is then fed into the user application as position and joint angle data (User Application), for example, to display a corresponding virtual hand in a virtual environment.

**Fig. 1.**
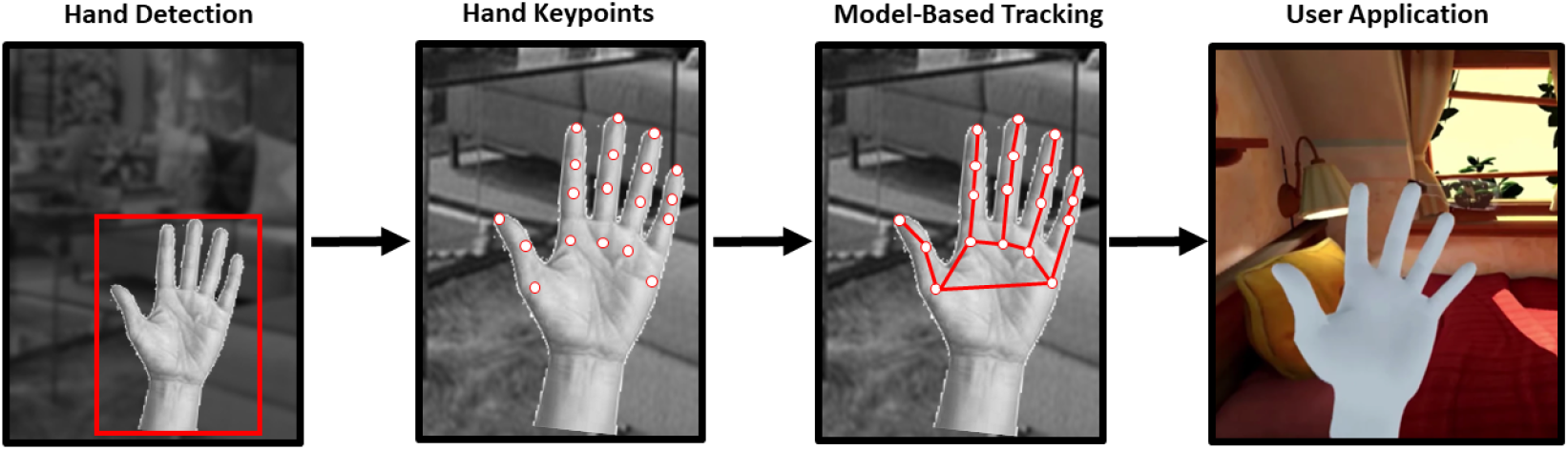
Meta Quest 2 computer vision driven multi-stage hand-tracking, including hand detection, key-point identification and tracking stages

The use of XR marker-less motion-tracking systems is spreading rapidly amongst researchers from a wide range of scientific fields due to the relatively small number of hardware components involved, the well-established technology, and reduced costs. For example, in robotics, accurate human hand and finger tracking data are used to safely teach robots to grasp various objects and to perform complex interactions in a simulation environment, before being deployed in the real world, [8]. In the field of human-computer interaction, marker-less hand-tracking allows for research into more natural ways of interacting with virtual objects in immersive environments, [9]. While in psychology and neuroscience, optical marker-less tracking has the potential to enable research into a wide variety of areas, such as tool use, social interactions and rehabilitation in VR, [10, 11].

However, despite the large variety of current applications and the future potential of these hand-tracking systems, there is little information regarding their tracking performance, such as positional and angular accuracy and delay, [12]. To address this, we present a robust methodological framework based on marker-based (ground-truth) tracking to evaluate such systems. We use this framework to assess the Quest hand-tracking system, which is one of the most popular and widespread examples of this technology. First, we introduce the methodological framework and its application to test the hand-tracking performance of the Quest, before describing our results. Finally, we provide an error-correction technique based on lens distortion to correct the Quest’s positional errors, [13, 14].

## 2 Methods

### 2.1 Participants

8 healthy volunteers (3 female, ages 25-48 years) were recruited from the University of Birmingham. All participants had a full range of arm and hand movements with normal or corrected-to-normal vision. This group of participants covered a range of hand sizes, with a measured circumference around the hand, ranging from 17.7*cm* to 22.2*cm*. Ethical approval was obtained through the University of Birmingham Psychology ethics panel.

### 2.2 Hardware

An Meta Quest 2 (Meta, formerly Facebook, 2022, build version 33.0) head-mounted-display (HMD) was rigidly held in a fixed position directly above the participant’s head but not blocking vision, with a downwards angle of 35^*o*^ towards a table with 18 equally spaced infrared (IR) reflective marker targets of 2*cm* radius. The 18 targets were glued in a 3 rows (A,B,C) and 6 numbered columns (1-6) to a 120*cm* x 55*cm* x 1.5*cm* piece of acrylic sheet, Figure 2. The grid of markers had a uniform spacing of 20*cm* in the three rows; the rows were 10*cm* apart and staggered to allow uniform sampling of the area within a rectangle of 110*cmx*20*cm*. The target panel was supported by an height-adjustable table at the center of the motion capture space, and was used at three heights from the HMD, low (80*cm*), mid (66*cm*) and high (52*cm*), such that the volume sampled covered 0.528*m*^3^, Figure 2. A VR-ready Windows 10 laptop with a dedicated Nvidia GTX-1060 graphics card served as our experimental control machine connecting to the motion capture cameras and the Quest. The full capture space was illuminated evenly from all sides.

**Fig. 2.**
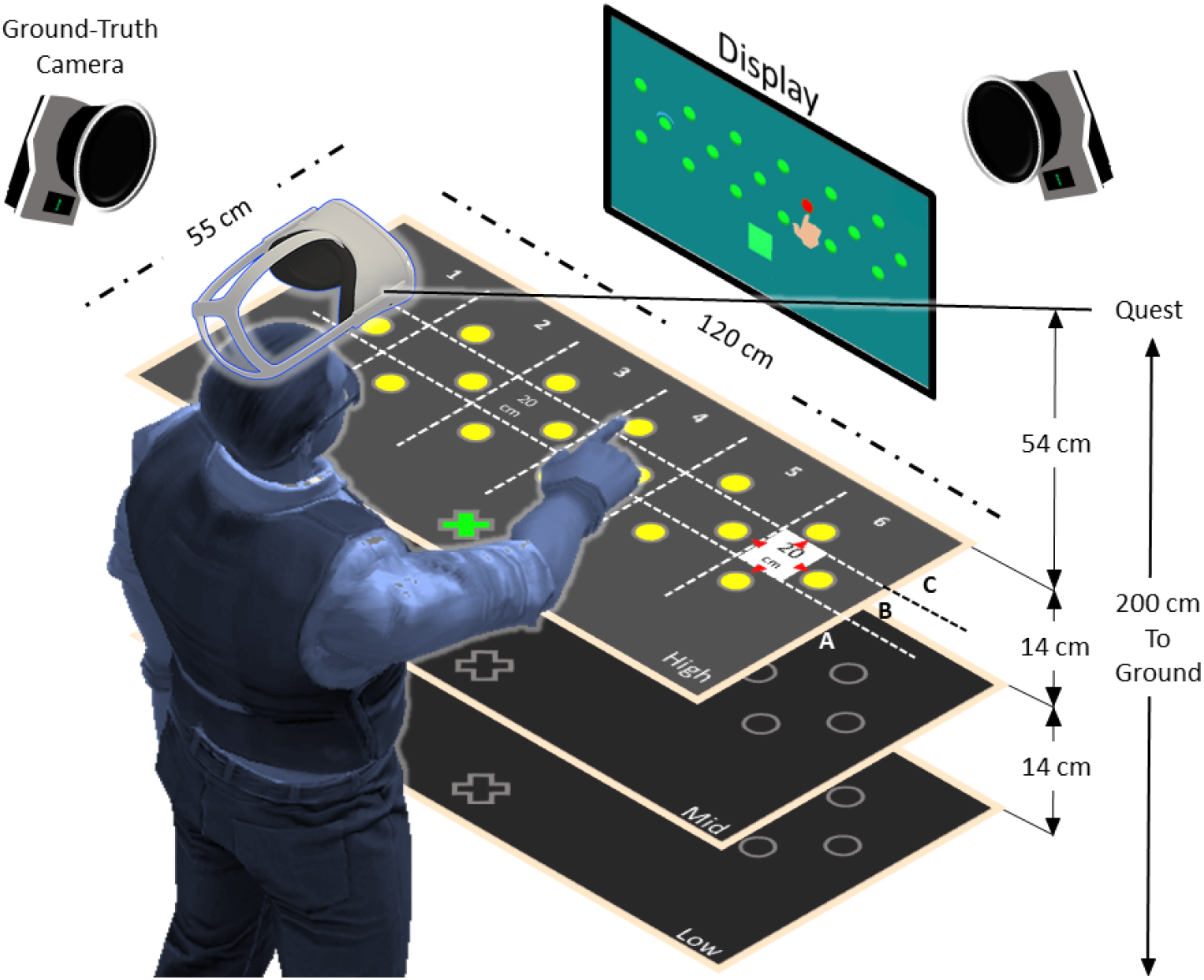
Experimental setup with the target panel, display, ground-truth cameras and user standing under the Meta Quest 2 HMD. The targets (yellow circles) are separated in rows A, B and C and columns 1 to 6 with a total of 18 targets per height. The green cross is the starting position

A set of 12 Oqus-350 cameras (Qualisys, 2021) were mounted on the ceiling aligned to cover the full tracking volume of the experiment, used as our marker-based ground-truth measurement device. This setup achieves sub-millimeter positional accuracy (0.033*cm*) at a sampling rate of 240*Hz*, based on a calibration procedure performed before the experiment. Figure 3A shows how we arranged a set of four 0.2*cm* IR reflective markers on the participants hand and fingers to identify the individual markers on known anatomical finger locations via a computer hand model provided by the Qualisys software, Figure 3B. This served as the ground-truth measure of fingertip position and timing.

**Fig. 3.**
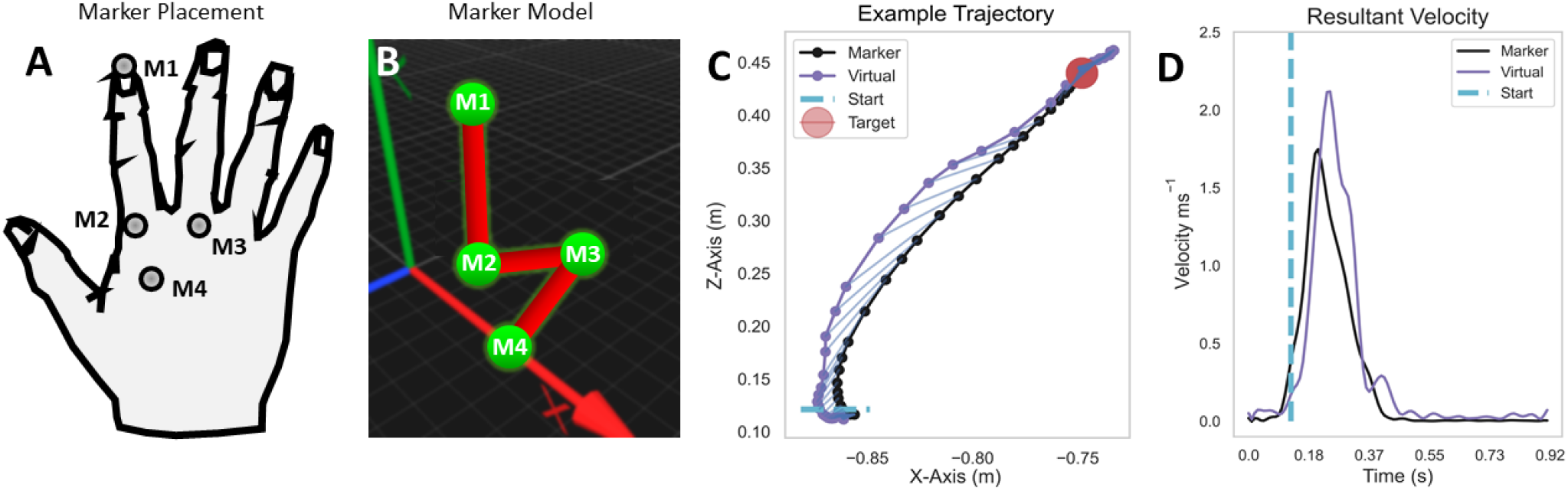
Hand Model. A: Infrared reflective markers (M1-M4) on the hand and fingers. B: Corresponding computer model of the hand and fingers from Qualisys ground-truth tracking software (perspective view). C: An example fingertip trajectory from the ground-truth marker and the Quest fingertip; D: The corresponding marker and virtual fingertip velocities

### 2.3 Software

A custom-built environment within The Unity game engine (Unity Technologies, *v*2021.1.9.*f*1) performed data recording and controlled the experiment progress, allowing hand and finger tracking data to be recorded from both ground-truth and Quest hand-tracking systems, synchronously at a sampling rate of 120*Hz*. The Unity-Oculus integration package and the Unity-Qualisys plugin allowed both systems to run from the same, custom-built, Unity environment. To use Quest native hand-tracking on the laptop we used a dedicated Oculus Link cable (Meta, 2022), which provided full access to all Quest functionality. We captured and stored the position of 18×3 target locations at the beginning of the experiment and transformed the right hand virtual index fingertip provided by the Quest and the index fingertip marker position data from the ground-truth system. This transform aligned the coordinate frames of both tracking systems using a simple target reaching procedure, where participants reached for the central target and the two most lateral target locations on the board at the beginning of each condition.

### 2.4 Experiments

To measure positional accuracy and angular performance two tasks were performed, a target reaching and a hand opening-and-closing task. These tasks covered two of the most common motor control scenarios, which require hand position and orientation data as well as finger joint angles.

#### 2.4.1 Target Reaching

This task involved participants standing in front of the target panel and reaching to one of the 18 targets of the current height in a random sequence. Each trial involved: 1. Place the right index finger on the starting position, indicated by a sphere on the target panel closest to the participant, see green cross in Figure 2A; 2. Reach to the highlighted target as indicated by a visual cue on the screen and by an auditory cue; 3. Maintain fingertip position on the target for 1 second; 4. Return to the starting position. Each target was repeated four times in each of the three blocked heights (low, medium and high) and three tempos 80, 120 and 160 beats-per-minute (bpm), resulting in 648 trials per height in total.

For each of the three tempos the metronome dictated how fast the participant should move between the staring position and the target for the target reaching task or how fast they should open and close their hand in the hand opening-and-closing task based on the temporal interval between the beats. Participants had the opportunity to listen to several beats before executing a trial.

#### 2.4.2 Hand Opening and Closing

Participants made a series of hand opening and closing movements for two minutes following one of the three tempos of 80, 120 and 160*bpm* while standing directly under the Quest, with the right hand raised chin-high and the palm rotated at a 45^*o*^ angle towards their face for optimal hand and finger tracking of the Quest device. One trial of continuous opening-and-closing movements per tempo was recorded lasting 60*s* each.

### 2.5 Data Processing

Each trial recorded marker positions on the index finger and hand both from the ground-truth system and the Quest for this analysis. Figure 3C shows an example fingertip trajectory from the target reaching task together with it’s corresponding velocity, Figure 3D. The dashed blue line in both figures indicates at which position and at what time point the fingertip leaves the starting region, following the start cue.

The following metrics were extracted for trials in the target reaching task: maximum fingertip velocity, positional accuracy (i.e. error) and path offset between the real and virtual fingertips. Time delay (lag) between the virtual and real fingertips were computed for both the target reaching and hand opening-and-closing tasks.

Next we describe how each metric was computed in more detail.

#### 2.5.1 Delay

Time delay was computed via the generalised cross-correlation tool using the positional and angular data from both Quest and ground-truth sensors, [15, 16]. We validated the delay results by randomly sampling from several trials using the separation between the two velocity peaks in time as number of samples between the fingertip velocity and angular velocity peaks of the two sensors, through a peak detection tool, [17], see the two red crosses in Figure 4B. To convert the number of samples back into a temporal delay (in milliseconds) the number of samples was multiplied by the sampling time, which is the inverse of the sampling rate:

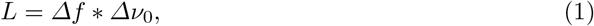

where *Δf* is the sampling time and *Δν*_0_ is the difference in the number of samples between the ground truth and Oculus Quest fingertip peak velocities.

**Fig. 4.**
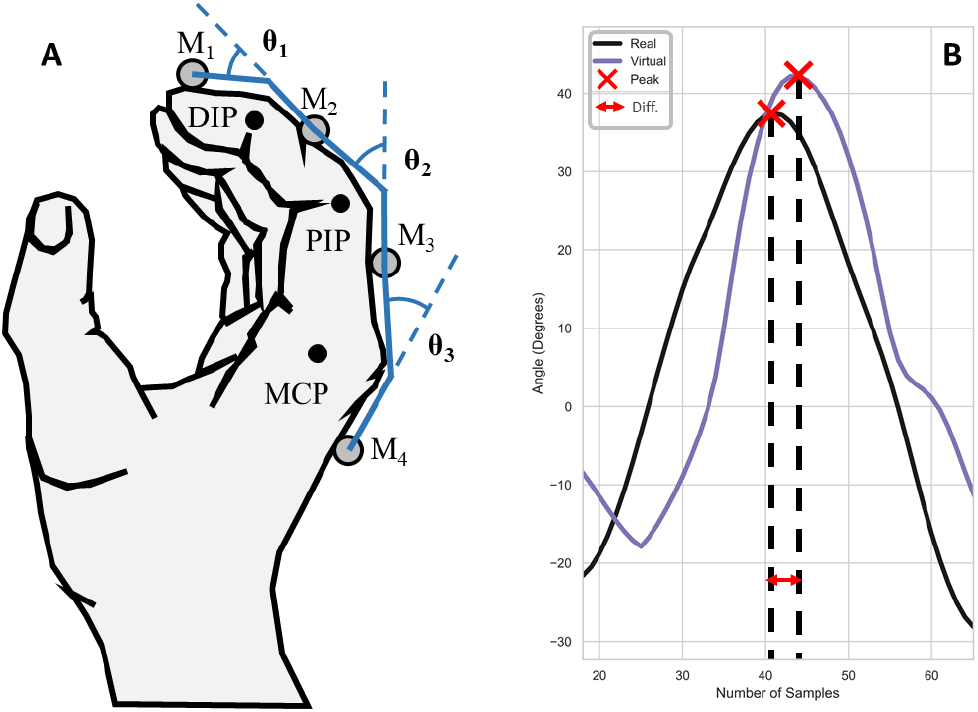
Hand opening-and-closing. A: Joint rotations showing the Distal (DIP), Proximal (PIP) and Metacarpo-phalangeal (MCP) joints of the index finger. B: Computing temporal delay between the virtual and real MCP joint angles using the peak angle (red crosses) as a metric for temporal alignment

#### 2.5.2 Static Positional Accuracy

This metric was designed to estimate how the static positional accuracy of the Quest changed as a function of distance from the hand, without any consideration of movement speed or delay. To this end, we computed the Euclidean distance using the last 30 samples of Quest fingertip and ground-truth marker positions, following a one second rest on the target to mitigate any temporal effects.

#### 2.5.3 Path Offset

To investigate the influence of movement speed i.e. the three tempo conditions, we computed a path offset metric, which was based on the Manhattan distance (D), equation 2. The distance between each sample was defined as the sum of the absolute differences between two vectors, in our case the trajectories of the ground-truth and Quest fingertip:

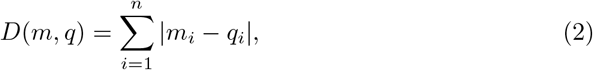

where (m,q) are the trajectory vectors for the ground-truth and Quest fingertip data, respectively.

To mitigate the effects of delay between the two tracking systems on this offset metric, the analysis shifted the Quest fingertip trajectory back in time by the average delay amount of 38.0 ms, as explained in section 2.5.1.

#### 2.5.4 Angular Error

For the hand opening-and-closing task the Quest provided joint angles directly, which we compared to the joint angles computed from the ground-truth markers. As a result, we had to change the marker arrangement to a row along the index finger. Figure 4A shows the hand during the opening-and-closing cycle, where the three joint rotation angles between the four phalanges of the index finger are illustrated based on the four markers (M1-M4). The inverse of the cosine function computes the angle between each pair of adjacent markers, M1-M2, M2-M3 and M3-M4, resulting in three angular values for each of the three joints: Distal (DIP), Proximal (PIP) and Metacarpo-phalangeal (MCP) of the index finger.

The time derivative of the joint rotations provides another metric, angular velocity error, which is computed from the average of the absolute difference between the real and virtual joint angle velocities for each of the joints, Figure 4B. This metric is delay corrected, meaning the error computations takes into account the delay in angular velocity between the Quest fingertip and ground-truth marker by shifting the Quest data back in time by the computed delay.

#### 2.5.5 Statistics

For each outcome metric, we used the R-Studio programming language (R-Studio v1.1456, R v3.61) together with the “lme4” statistics package to fit a generalized linear mixed-effects model (GLMM) using the three fixed effects (tempo, height and target location) and subject as a random factor. The default logistic regression (logit) link function was used in all applications of the GLMM function using random intercepts. To report the resultant GLMM model the “Anova” function from the “car” statistics package was used, which provided the F-value, degrees of freedom and p-values. For post-hoc analysis we applied a pairwise least-squares means comparisons from the “multcomp” and “lsmeans” R statistics packages with sequential Bonferroni adjustments.We chose the GLMM model to account for non-normally distributed data, as was present in a pilot data set. Most metrics showed a beta distribution.

## 3 Results

### 3.1 Delay

During the reaching task, the Quest hand and finger tracking data showed an average delay of *M* = 38.0 *ms, SD* = 31.9 *ms* when compared to the ground-truth data. The GLMM showed that the delay was independent of height *F*(2, 2478.9) = 1.1, *p* = 0.16, tempo *F*(2, 22477.8) = 1.3, *p* = 0.08 and target location *F*(17, 2477.1) = 11.5, *p* = 0.43. This mean delay was used to temporally align all trajectories in subsequent analyses.

### 3.2 Static Positional Accuracy

This error metric showed a characteristic pattern with larger static positional errors in the periphery of the Quest’s visual field, i.e. when the fingertip was farthest away and to periphery of the HMD, Figure 5. In contrast, height did not have a significant effect on positional accuracy, *F*(2, 2477.5) = 1.3, *p* = 0.27. For this metric we do not consider tempo as a factor, because we were only interested in static positional accuracy. Therefore, positional error was averaged across the three heights and tempos for the subsequent analysis. The GLMM showed a main-effect for planar target location on positional accuracy, *F*(17, 2477.1) = 17.4, *p <* 0.001.

**Fig. 5.**
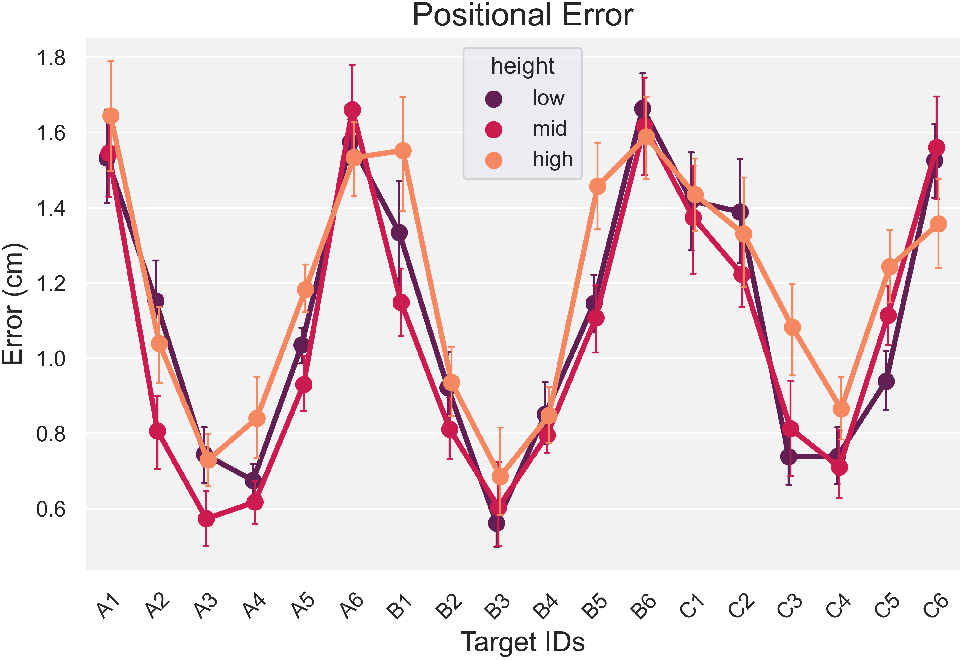
Positional error across the three heights and 18 targets, averaged across the three tempos. Error bars represent standard error of the mean

Specifically, a post-hoc analysis showed peripheral targets (see Figure 2 for target locations), i.e. A1, A6, B1, B6 and C1, C6, had a significantly higher positional error (*M* = 1.3*cm, SD* = 0.46*cm*) compared to the positional errors measured for the central targets (targets 2−5 for rows *A*−*C*), (*M* = 0.79*cm, SD* = 0.39*cm*), (*t* = 7.2, *p <* 0.01, 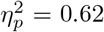). There was a significant interaction between height and target, *F*(34, 2477.1) = 2.3, *p <* 0.001. A subsequent post-hoc analysis on this interaction showed the error for central-mid-level targets (*M* = 0.75*cm, SD* = 0.1*cm*) was significantly smaller, (*t* = 8.2, *p <* 0.001, 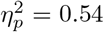), compared to peripheral targets at the low and high level (*M* = 1.3*cm, SD* = 0.1*cm*).

### 3.3 Path Offset : Trajectory Error

This error metric compared the Quest and ground-truth fingertip trajectories during movement from the starting position to the target on a sample-by-sample basis (i.e. error across a movement). As a result, the baseline delay between the two sensors was corrected by shifting the Quest data samples forward by 38.0 *ms* to align with the ground-truth data. This step ensured the measurements were taken between temporally synchronous samples and therefore better represented path offset without contamination with delay. As shown in Figure 6A, path offset was heavily influenced by the speed of the movement and the relative distance to the HMD. Specifically, a GLMM revealed a significant main effect for tempo (*F*(2, 2478.0) = 60.9, *p <* 0.001), height (*F*(2, 2479.2) = 11.8, *p <* 0.001) and target (*F*(17, 2477.1) = 7.8, *p <* 0.001). In addition, there was a significant interaction between tempo and height (*F*(4, 2477.6) = 6.2, *p <* 0.001) and marginal interactions between tempo and target (*F*(34, 2477.0) = 1.4, *p* = 0.054) and height and target (*F*(34, 2477.1) = 1.5, *p* = 0.057). Post-hoc tests showed that the mid-layer height had a significantly lower path offset (less error), *t* = 2.2, *p <* 0.05, 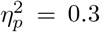, compared to the other heights in the 120*bpm* (*M* = 4.1*cm, SD* = 1.2*cm*) and 160*bpm* (*M* = 4.3*cm, SD* = 1.2*cm*) tempo conditions but not for the 80 bpm condition, Figure 6B.

**Fig. 6.**
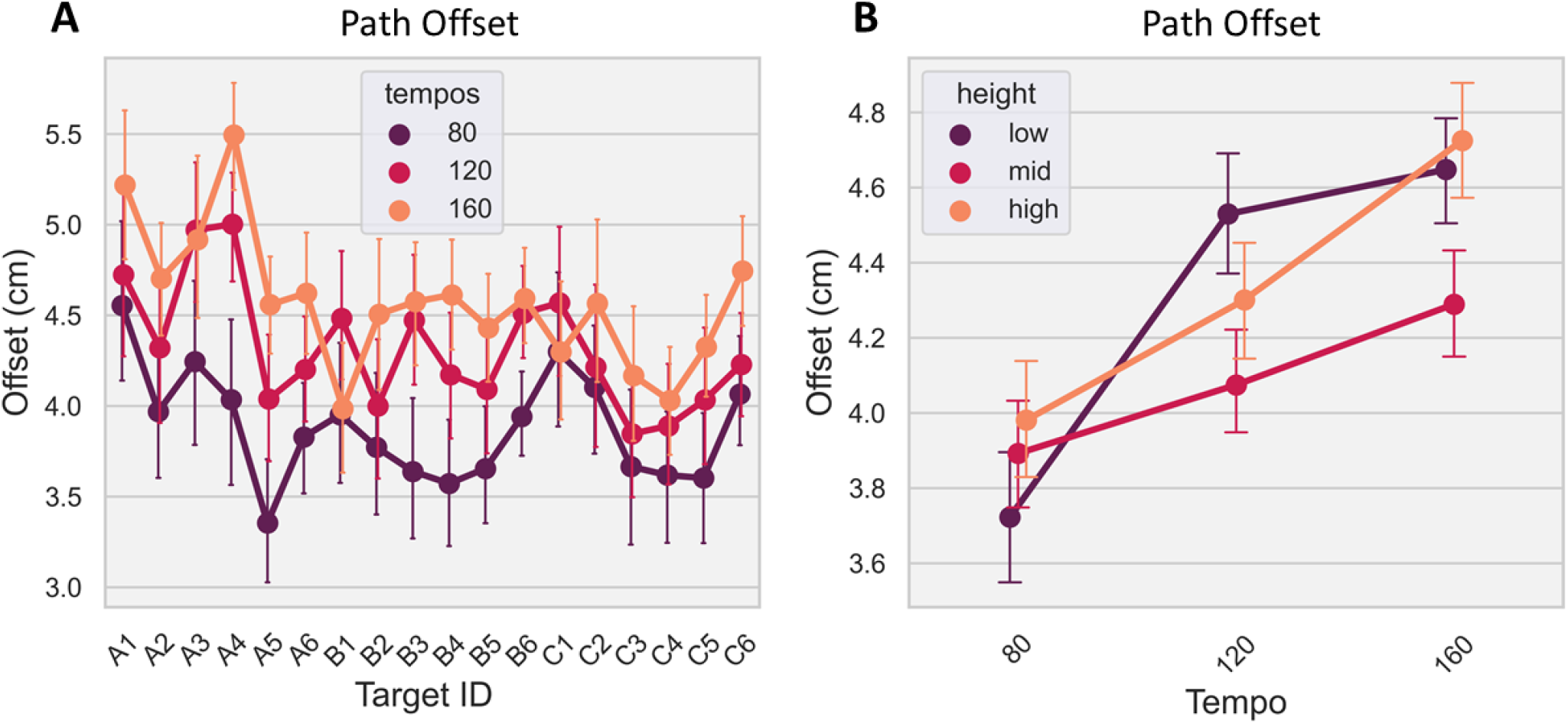
A: Path offset across the three tempos and 18 targets, averaged across the three heights; B: Path offset with standard error across the three tempos and heights, averaged across the targets. Error bars represent standard error of the mean.

### 3.4 Angular Error

The hand opening-and-closing task provided joint angular rotations for each pair of adjacent joints of the right hand index finger via the ground-truth markers, figure 4A. We compared these angular estimates with the joint angle measures provided directly by the Quest hand tracking, including angular velocity error. Figure 7A shows angular error from the Quest changed with increasing rotation speed across all three joints, with error increasing for the MCP joint from 11.5^*o*^ at the 80*bpm* level to 16.7^*o*^ at the 160*bpm* tempo. In contrast, error decreased in the other two joints with increasing tempo. The GLMM analysis confirmed these observations with a main-effect of tempo *F*(1, 893.0) = 47.8, *p <* 0.001 and joint *F*(2, 893.0) = 15.5, *p <* 0.001. There was also a significant interaction between tempo and joint, *F*(2, 893.0) = 49.8, *p <* 0.001. A post-hoc test showed interactions between these two factors. Angular error between the three joints at the 80*bpm* tempo level was not significantly different, (*t* = 1.1, *p* = 0.95, 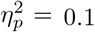). However, for the 120*bpm* and 160*bpm* tempos, angular error was significantly different between the three joints, (*t* = 10.4, *p <* 0.001, 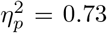), confirming the Quest’s joint angle estimates increased in error with faster rotational movements in the MCP joint. The overall average angular error across the three tempos and joints was *M* = 9.6^*o*^, *SD* = 6.2^*o*^.

**Fig. 7.**
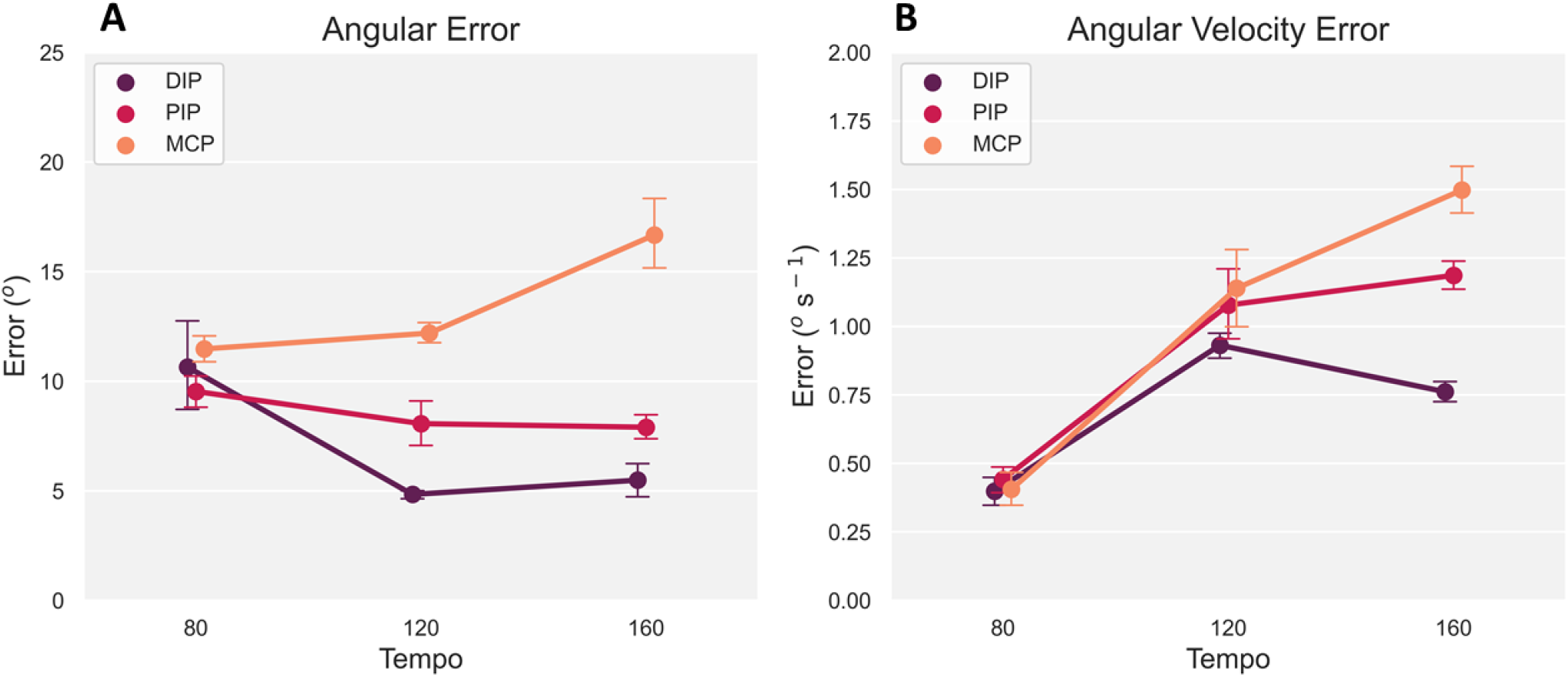
A: Angular error with standard error between ground-truth marker and Meta Quest averaged across the three joints (DIP, PIP and MCP). B: Corresponding angular velocity error with standard error between the ground-truth marker and Meta Quest sensors

Angular velocity error presented an increasing trend across the three tempos showing larger separation in error with faster rotational movements of each joint, Figure 7B. A GLMM analysis confirmed this trend for tempo, with a main-effect, *F*(2, 890.0) = 43.0, *p <* 0.001. The three joints were not significantly different in velocity error from one another, *F*(2, 890.0) = 0.32, *p* = 0.73. However, there was a significant interaction between tempo and joint, *F*(4, 890.0) = 20.5, *p <* 0.001. A post-hoc analysis showed that the separation in velocity error between the PIP and DIP joints at the 160*bpm* tempo was significant, (*t* = 5.3, *p <* 0.001, 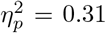). This significant interaction is also true for the other two joint pairs at the 160*bpm* tempo level, (*t* = 12.5, *p <* 0.001, 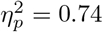).

### 3.5 Error Correction

The aim of this section is to present position-correction results from the previously-identified errors. Figure 8 illustrates the Quest’s static positional errors across it’s entire tracking volume, with accuracy worsening in the periphery of the visual field. This seems especially true for the lowest height i.e. the furthest distance between the target panel and the Quest HMD (bottom row in Figure 8).

**Fig. 8.**
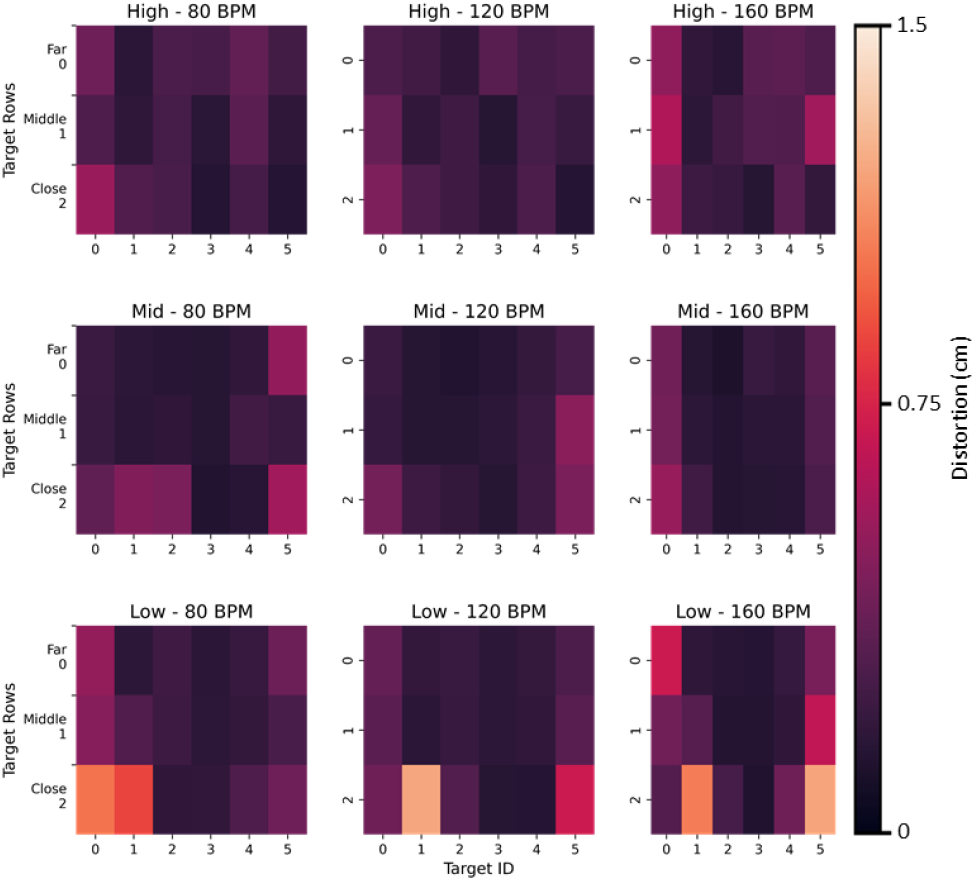
Heat-maps of the average positional error for a particular tempo and height across all participants. Each row of plots is for a particular target-panel height starting with the highest to lowest heights (top to bottom) and each columns represents one of the three tempos (80, 120 and 160*bpm*) from left to right

Together with the ground-truth data, these error matrices were used to construct a lens distortion model. To make an analogy, distortions in images taken by a camera are caused by imperfection in the camera’s lens, and to correct for these distortions we would need to characterise and reverse the offset at each location on the image. For a subset of the positional error data (80 per cent), we defined a fifth-order polynomial (*N* = 5) lens distortion model (equation 3) and applied this to each of the three tempos and heights, with the four-trial average Quest and ground-truth positions (i.e. distorted and un-distorted positions, respectively) as input to the model:

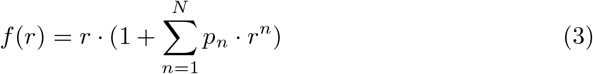

where r is the raw positional data (e.g. decimal number representing position in z axis) and *p*_*n*_ is the *n*^*th*^ parameter.

Based on this input, a least-squares optimisation tool, [17], computed a set of five lens correction parameters by minimizing the error between the Quest’s raw input and the ground-truth positions through an iterative optimisation procedure, Table 1. A paired t-test showed these five parameters were sufficient to improve the positional error of the remaining 20 per cent of unseen raw position data, as this resulted in corrected position data that was significantly closer to the ground-truth (*t*(3.1), *p <* 0.001) as opposed to the raw Quest positions, Figure 9.

**Table 1.**
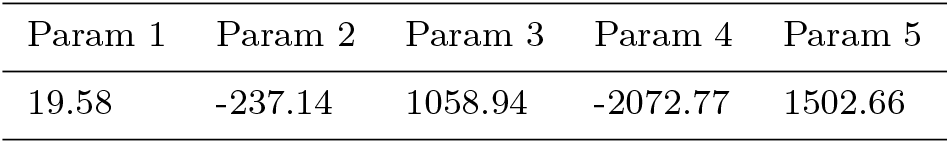
Lens distortion correction parameters for reconstructing the Meta Quest 2 positional tracking errors

| Param 1 | Param 2 | Param 3 | Param 4 | Param 5 |
| --- | --- | --- | --- | --- |
| 19.58 | -237.14 | 1058.94 | -2072.77 | 1502.66 |

**Fig. 9.**
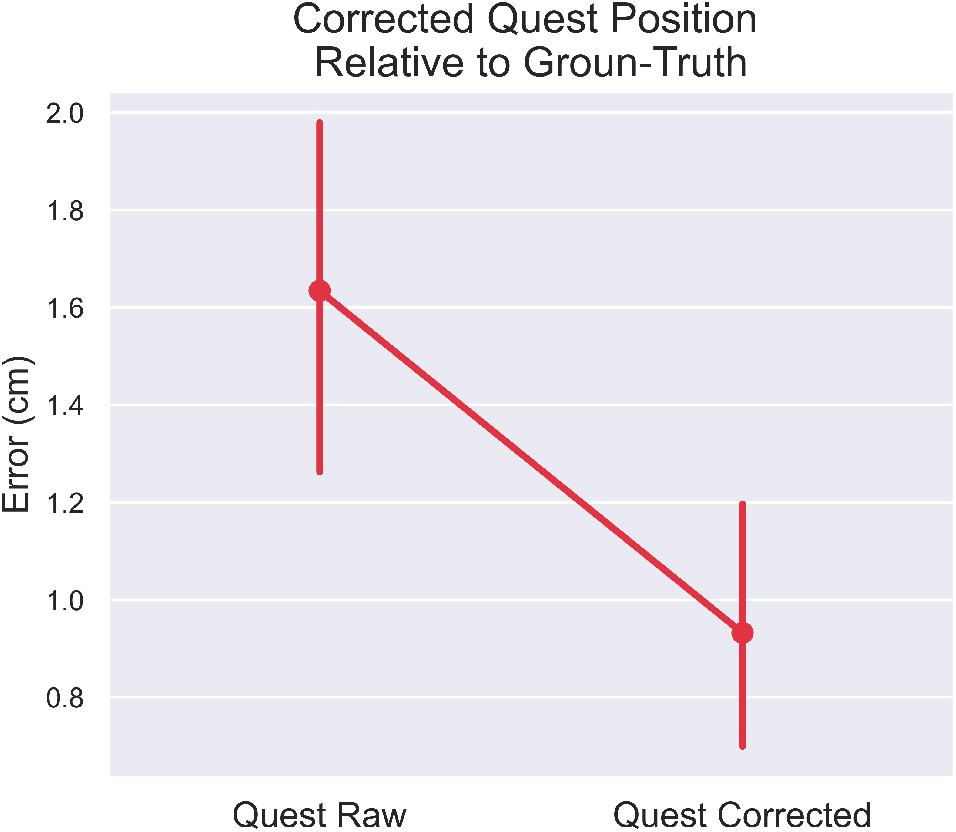
Correcting Meta Quest 2 tracking error with standard error using five lens correction parameters and a polynomial lens distortion model for target reaching across a large (120 cm x 55 cm x 80 cm) tracking space. This set of data was not used when the parameters for the lens-correction model were computed

## 4 Discussion

This article presents a novel methodological framework to assess the accuracy of VR optical hand and finger tracking systems, such as the popular Meta Quest 2 VR headset. Furthermore, we present a novel approach to account and correct for tracking errors based on a lens distortion model. The results show a set of useful metrics such as accuracy (static measure of positional error), path offset (trajectory error during motion), angular error and temporal delay. We believe these metrics could be highly informative when deciding whether to use such hand tracking systems within environments which require high spatial and temporal accuracy (i.e. motor control/learning research).

The method presented here provides clear quantitative evidence to show that static positional errors are largely affected by the distance between the hands and fingers from the central view area of the headset, with larger positional errors further away from the center. We speculate this is most likely due to a combination of factors related to the headset’s camera lens distortion correction model and the camera’s resolution. Fingers in the periphery of vision are distorted to a larger degree compared to the center, resulting in the Quest’s hand tracking model to underestimate the finger’s true position. Future work could focus the analysis on the directionality of the positional errors, in order to confirm that peripheral position estimates are indeed overestimated by the model compared to the central position estimates.

For path offset, height, tempo and target location all had a significant impact on this cumulative metric, because in any given trial hand motion could cover a large tracking space (i.e. nearer or further away from the headset), and with one of three motion speeds. Faster movements resulted in larger offsets. In this case, it is likely the Quest’s pre-trained machine-learning model receives fewer frames of the hand in rapid motion, such as in the 160*bpm* condition. Compared to the 80*bpm* condition, these fewer frames of the hand could result in fewer data points for the algorithm to accurately estimate key-points around the hand and fingers [4].

However, because the Quest uses a proprietary hand-tracking algorithm, it is difficult to know for certain which computational stage would be most affected by rapid hand movements. Google’s open-source, camera-based MediaPipe hand-tracking algorithm provides an insight into the inner-workings of the Quest hand-tracking algorithm, who show that their machine-learning algorithm is based on a set of assumptions about skin tones, textures, hand sizes, camera model and hand movement parameters, [18]. Based on this, it is fair to assume that the Quest’s pre-trained algorithm is also based on a similar set of assumptions. An equally viable argument for why tracking performance decreases during fast motions could be made based on the other stages of the algorithm, such as the hand detection or application of inverse kinematics stages. In our case, another impact-full test parameter could be related to skin tones. Although our design covered a relative diverse set of hand sizes, we only recruited participants with skin tones in the light range. This is an important area to consider when planning to use the Quest hand tracking for research related purposes.

The overall average angular error was 9.6^*o*^, which was differentially affected for each joint with increasing tempo. We did find that faster rotational movements of the MCP joint significantly increased angular errors. This could be due to self-occlusions, where the other finger phalanges block direct visual line of sight of the MCP joint when closing the hand in a palm-facing direction. The MCP joint is occluded and the algorithm has to rely on previous joint estimates, which could be incorrect, or estimate the occluded joint angle based on other parameters, such as the other visible joints and the overall hand pose, all of which would be a less accuracy estimate of the actual joint angle. However, this speculation requires further testing with a new experimental design, such as performing hand opening-closing at different viewing angles relative to the Quest’s cameras.

Angular velocity error was significantly lower in the low speed condition (80*bpm*) compared to the 120*bpm* tempo but not the 160*bpm* tempo condition. One explanation for this is self-occlusions of the proximal joint by the back of the hand when making a fist during the hand opening-and-closing task. This interrupts the stream of images of the hand to the algorithm, which pushes the estimates to rely on a sparser data set to estimate joint angles, resulting in larger errors. To test this hypothesis, more hand opening-and-closing task data are required at different viewing angles relative to the headset so that self-occlusions can be avoided.

Delay is one of the most important metrics in gaming, research and robotics because it determines the limits in applicability. The results showed an average delay of 38.0*ms* in the Quest hand-tracking data, which was unaffected by motion speed or location of the hand relative to the headset. Therefore, this delay is most likely caused by the headset’s optical sensors, processor and wireless communications capabilities. As this is a hard limit, the temporal delay must be taken into account when using the Quest within any environment in which the tracking data is being used to assess motor performance.

To expand the usable range of the Quest hand-tracking for static positional estimates, we applied a lens distortion model across the entire tracking space to correct for errors. With only five parameters, it was possible to correct for positional errors across the Quest’s tracking volume. Whilst this appears to be be a powerful technique to correct for the spatial errors observed in the hand tracking data, further analysis is required to test the limits of this approach. For example, at what level of tracking error does the reconstruction break down by providing worse results than the raw tracking data from the optical machine-vision system?

Despite the positional errors and the presence of lag, there is little reporting of clear discrepancies between actual and virtual hand movements within VR apps that use marker-less hand tracking. This indicates that within an immersive environment these discrepancies may not be noticeable. However, when deciding whether to use these hand tracking systems for experimental paradigms, researchers should be aware of these errors and evaluate the consequences for their research goals. For example, in the field of motor learning, small measurement errors may be highly undesirable. However, if the emphasis is on rehabilitation, and encouraging the use of the hands, then the consequences of accurate measurement may not be of such importance, in comparison to the immersion in the task. Similarly, research focusing on social interaction may be less affected than tasks involving object manipulation. The results presented here enable researchers to make informed choices and also follow our methodology to correct for some part of these errors. Our results also enable certain recommendations when planning experiments or tasks, such as keeping the hands nearer the centre of the tracked volume directly in front of the HMD and emphasising postures which do not occlude the fingers. Furthermore, given the sensitivity of some metrics to movement speed, the type of movements being studied should be carefully chosen. The framework detailed here also stands as a repeatable methodology to measure hand tracking accuracy of other devices and future generations of XR HMDs.

In summary, this article provides a robust methodological approach for assessing the temporal and spatial accuracy of VR-based marker-less hand-tracking systems. Analysis of the Meta Quest 2 indicated clear temporal and spatial limits of the device for tracking hand-based movements which either need to be adjusted for or taken into account when making conclusions based on the data.

## Acknowledgement

This work was supported by the European Research Council Starting Grant (MotMotLearn 637488) and European Research Council Proof of Concept Grant (ImpHandRehab 872082). We would like to thank Prof. Alan Wing, Mohamed Maaroufi and the SyMoN lab for supporting this project.

